# Muscle function during locomotion on skis at varying speed and incline conditions

**DOI:** 10.1101/2022.12.27.522016

**Authors:** Amelie Werkhausen, Anders Lundervold, Øyvind Gløersen

## Abstract

The human musculoskeletal system is well adapted to use energy efficient muscle-tendon mechanics during walking and running but muscle behaviour during on-snow locomotion is unknown. Therefore, we examined muscle and muscle-tendon unit behaviour during diagonal style roller skiing at three speed and incline conditions.

We assessed lower leg muscle and muscle-tendon unit mechanics and muscle activity in thirteen high-level skiers during treadmill roller skiing using synchronised ultrasound, motion capture, electromyography and ski-binding force measurements. Participants skied using diagonal style at 2.5 and 3.5 m·s^−1^ at 5°, and at 2.5 m·s^−1^ at 10°.

We found an uncoupling of muscle and joint behaviour during most parts of the propulsive kick phase in all conditions (*P*<0.01). Gastrocnemius muscle fascicles actively shortened ~9 mm during the kick phase, while the muscle-tendon unit went through a stretch-shortening cycle. Peak muscle-tendon unit shortening velocity was five times higher than fascicle velocity (375 vs 74 mm·s^−1^, *P*<0.01). Increased incline was met by greater muscle activity (24%, *P*=0.04) and slower fascicle shortening velocities (34 vs. 45 mm·s^−1^, *P*<0.01). Increased speed was met by greater peak muscle activity (23%, *P*<0.01) and no change in fascicle shortening velocity.

Our data show that muscle behaviour was uncoupled from the joint movement, which enables beneficial contractile conditions and energy utilisation during diagonal style at different slopes and speeds. Active preloading in the end of the glide phase may benefit the mechanisms.

**Summary statement:** We examined muscle function during diagonal style cross country skiing in competitive cross-country skiers. Our data show an uncoupling of muscle and joint behaviour in the lower leg during skiing.

## Introduction

The human musculoskeletal system is well adapted to its two primary forms of terrestrial locomotion; walking and running (e.g. Arnold and Delp, 2011; Farris and Sawicki, 2012). Both gaits are characterized by ground contact periods where the foot is stationary on the ground, held in place by friction forces. During the ground contact phases of these gaits, elastic energy is stored and released as the muscle tendon unit (MTU) elongates and shortens, allowing leg muscles to operate at substantially lower velocities than the MTU (Ishikawa et al., 2005; Lichtwark et al., 2007; Lichtwark and Wilson, 2006). Utilisation of elastic energy and the energy-efficient way to generate muscle tension explains why these gaits are metabolically efficient (Fletcher and MacIntosh, 2015). However, this energy efficient contraction mode is not apparent in all locomotor activities. Recent studies examining muscle behaviour during race walking and rowing, showed that fascicles followed the stretch-shortening cycle of the MTU, resulting in less efficient contractile patterns (Cronin et al., 2016; Held et al., 2022).

Diagonal stride (DS) is a skiing gait typically used in uphill terrain. It is a four-limbed gait, where arms and legs move in opposite phase (i.e. stance on left ski and right pole simultaneously, and vica versa). The arms are in contact with the ground through ski poles, but they only carry a small fraction of the load compared to the legs (7-8% of the body weight on average over a cycle, Vähäsöyrinki et al. (2008)). Similar to running, DS has in-phase fluctuations of kinetic- and gravitational energy (Pellegrini et al., 2014). It has therefore been suggested to be a “bouncing gait”, similar to running, which utilizes the MTUs capacity to store elastic energy. However, DS lacks the aerial phase seen in running. Instead of an aerial phase, DS has a glide phase, where the skier glides on one ski as the contralateral leg is in its swing-phase. Hence, DS has been characterized as a form of “grounded running” (Pellegrini et al., 2014). The energy-efficient contraction dynamics seen in running can also appear in these bouncing gaits without aerial phases (i.e., grounded running; Davis et al. (2020); Reinhardt and Blickhan (2014)). However, by analysing fluctuations of mechanical energy, Kehler et al. (2014) concluded that the mechanical energy fluctuations in DS are most likely dissipated as frictional heat during the glide phase, indicating that the gait might not take advantage of the energy-saving mechanisms of other bouncing gaits. Hence, the term “grounded running” might be an inappropriate description for the gait. Nonetheless, joint-level kinematic and kinetic analyses of diagonal stride have shown that the lower leg muscle-tendon units undergo a stretch-shortening cycle during the kick-phase (Komi and Norman, 1987; Vähäsöyrinki et al., 2008). This suggests that the last part of the glide phase is used to produce pre-tension, enabling elastic tissue to stretch by active muscle contraction. A pre-tension during the glide phase could enable similar muscle-tendon behaviour as in jumping from a still position, where a catapult like function of the lower leg was observed (Farris et al., 2016; Kurokawa et al., 2001). The stretch-shortening cycle of the MTU described by Komi and Norman (1987) is in line with this reasoning. However, their study did not distinguish between muscle- and tendon-contributions to joint movement, which hindered conclusions on the muscles’ contraction mode. Hence, it is still unclear if, and how, the diagonal stride technique can leverage energy-saving mechanisms found in bouncing gaits.

In running, muscle contraction velocity is relatively unaffected by changes in speed, incline, or external load (i.e. vertical force). Rather, the main adaptations to increased demand (i.e. speed, incline, or vertical load) is increased muscle activation (Farris and Sawicki, 2012; Lichtwark and Wilson, 2006). For DS, the kick-phase duration increases with incline and decreases with speed (Nilsson et al., 2004; Pellegrini et al., 2013). If the pre-tension during the late glide- and early kick-phase of DS is initiated by active muscle contraction, it is likely that fascicle contraction velocity and muscle activation will increase with speed, since less time is available for the kick-phase. Muscle activation is also likely to increase with incline, as the resisting force increases. Fascicle contraction velocity, however, might decrease with increasing incline, as the kick-phase duration becomes longer.

The aim this study was to explore gastrocnemius medialis muscle and muscle-tendon unit behaviour during diagonal stride roller skiing, and to investigate the effect of speed and incline on these variables. We hypothesised, active shortening of the gastrocnemius muscle, while the MTU undergoes a stretch-shortening cycle during the late glide-phase and kick-phase. Furthermore, with increased speed, we expected greater muscle activation and higher fascicle shortening velocities during the kick-phase, due to a shorter kick duration. At greater incline, we expected greater muscle activation during the kick but similar fascicle velocity, due to increased kick duration when the resisting force is greater.

## Materials and Methods

### Participants

Thirteen XC-skiers (8 male and 5 female; age: 26.5 ± 4.8 years; height: 179.3 ± 8.0 cm; body mass: 72.8 ± 6.2 kg; and Fédération Internationale de Ski (FIS) ranking: 107.4 ± 62.7 points) with at least 5 years of competitive experience in XC-skiing at national or international level, gave written consent to participate in the study. All participants were familiar with the use of roller skis on a treadmill. The protocol was approved by the ethics committee of the Norwegian School of Sport Sciences (137-180620).

### Experimental protocol

Prior to testing, participants performed 10 minutes of low intensity roller skiing on a motorised treadmill used during the whole test session (Rodby, Vänge, Sweden) as warm-up and to ensure that the roller ski rubber wheels and bearings reached a proper temperature for testing (Ainegren et al., 2008). The roller skis’ (IDT Sport classic, Lena, Norway) back and front wheels were interchanged, so that the ratchet wheel mechanics was mounted in the front wheel. The participants used ski poles (Swix Triac 3.0, BRAV Norge, Norway) with special ferrules for use on treadmill surfaces and self-selected length, within official international regulations (FIS, 2020; maximum 83% of body-height).

During the test, participants roller skied with diagonal stride technique at three different conditions with varying speed and incline: 2.5 m·s^−1^ and 5° incline (DIA_ref_), 3.5 m·s^−1^ and 5° incline (DIA_fast_) and 2.5 m·s^−1^ and 10° incline (DIA_steep_). Data were recorded twice for each condition for ten seconds and each trial lasted for ~30 s. Ultrasonography, surface electromyography (EMG), force sensitive resistors (FSRs) under the ski binding and 3D kinematics data were synchronized by a trigger signal. Participants were given 2 minutes of rest between each trial.

### Joint kinematics and muscle-tendon unit length

We used a 12-camera 3D motion capture system (Qualisys 400 and 700-series, Qualisys AB, Gothenburg, Sweden) to record the three-dimensional trajectories of 26 reflective markers (12 mm, Qualisys AB) at 150 Hz. Twelve markers were placed at key anatomical landmarks on the pelvis and the right leg. The pelvis segment and the joints centers of the hip were defined by markers on the right and left anterior-superior and posterior-superior iliac spine (Bell et al., 1989). The knee joint center was defined as the mid-distance between markers on the medial and lateral epicondyles, and ankle joint center was defined as the mid-point between the lateral and medial malleoli. To track the foot segment, markers were placed on the calcaneus and the first, second and fifth metatarsal head (these markers were placed on the ski boot, not directly on the skin). The movement of the thigh and shank were tracked with a three-marker cluster positioned midway along these segments. Ankle, knee and hip angles were calculated based on the model described by Robertson et al. (2013) (Visual 3D, C-motion, Germantown, MD, USA). Eight additional reflective markers were placed on relevant sites of the equipment (i.e., at the back and front wheel of each roller ski, and one 15 cm below the handle of each pole, and one 60 cm above the tip for each pole). Reference frames for all segments were defined during a static capture were participants stood still holding their arms straight out in front. Marker trajectories were filtered using a second order lowpass digital zero-phase Butterworth filter with a cut-off frequency 15 Hz.

We calculated individual gastrocnemius medialis muscle-tendon unit length based on ankle and knee joint angles along with shank segment length (Hawkins & Hull, 1990). Shank length was defined as the mean distance between the lateral femoral epicondyle and the lateral malleolus of the right leg. Muscle-tendon unit velocity was calculated by numerical differentiation of the length.

### Muscle fascicle length

Gastrocnemius medialis muscle fascicles were imaged by an ultrasound linear array transducer (LV8-5N60-A2 ArtUs, Telemed, Vilnius, Lithuania), which was placed over the muscle belly using a custom-made holder and self-adhesive tape. B-mode images were recorded at 117 Hz with a field of view of 50 mm and depth of 65 mm. To make sure the cable of the ultrasound system did not obstruct the participants, the ultrasound control unit was placed in a box above the participants with an adjustable pulley system and cables were fastened with tape. A semi-automated tracking algorithm was used to analyse fascicle length and pennation angle (Cronin et al., 2011; Farris & Lichtwark, 2016). Fascicle length was defined as linear trajectory between the superficial and deep aponeurosis in direction of the visible fascicle fragments. Linear extrapolation was used when fascicle insertion exceeded the ultrasound image.

Fascicle length and pennation angle were resampled to the motion capture system’s frequency (150 Hz) using linear interpolation and filtered using the same digital lowpass filter. Subsequently, fascicle velocity was calculated by numerical differentiation of the length.

### Forces under the ski bindings

We placed two force sensitive resistors (Flexiforce A201, Tekscan, load range 0-445 N, sampled at 2100 Hz) under the front and back of the ski binding to estimate vertical forces under the skiing boot. To ensure that all forces between foot and ski were transmitted through the FSRs, a custom-made rectangular disc with a circular piston (⌀ 10 mm) pressed against the FSR’s sensing areas. Forces were calibrated for each participant using three measurements: (1) unloaded right ski, (2) participant stood on the forefoot, and (3) participant stood on the heel. The calibration factors for the FSRs (gain and offset) were calculated by solving the system of equations:

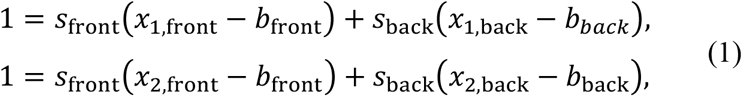

where *b*_front_ and *b*_back_ are the offset for each sensor (acquired separately from the unloaded recording), *x*_i,front_ and *x*_i,back_ are the FSR outputs from the forefoot (*i*=1) and heel (*i*=2) recordings, and *s*_front_ and *s*_back_ are the sensor gains. Force data were filtered with a second order lowpass digital zero-phase Butterworth filter with a cut-off of 30 Hz.

### Muscle activity

EMG data were recorded at 2100 Hz using a telemetered system (Aktos, Myon, Schwarzenberg, Switzerland). Bipolar surface electrodes (24 mm diameter; Kendall Arbo H124SG electrode, Coviden, Minneapolis, USA) were mounted over the muscle belly of gastrocnemius lateralis (GL), soleus (SO) and tibialis anterior (TA) as recommended in the Seniam guidelines (Hermens et al., 2000) after the skin was shaved and rinsed with isopropanol. EMG data were filtered with a second order bandpass Butterworth filter with passband from 10 to 500 Hz before a moving root mean square with a window length of 100 ms was calculated. RMS-filtered EMG data were individually normalised to its maximum value during DIA_ref_. The threshold to classify the muscle as active was set to 25% of peak based on the recommendations of Ozgünen et al. (2010).

### Cycle characteristics

A cycle was defined by the start of the kick until the start of the subsequent kick. The initiation of the kick phase was defined by the roller skies velocity becoming equal to that of the treadmill belt with a threshold of 0.5 km h^−1^, i.e., when the ski is no longer moving relative to the ground surface (Pellegrini et al., 2014). We used the reflective marker placed on the anterior wheel of the right roller ski as reference for the position of the ski in anterior-posterior direction. The skies velocity was calculated using numerical differentiation of the marker position. To calculate velocity along the treadmill band, we rotated the coordinate system to be parallel to two markers placed along the progression axis on the treadmill. The velocity data were filtered using a bidirectional second order Butterworth filter with a cut-off frequency of 20 Hz.

Each cycle was further divided into three phases: kick phase, swing phase, and glide phase. The start of swing phase was defined as when the skis velocity was greater than the treadmill velocity (with threshold of 0.5 km h^−1)^. The start of the glide phase was defined as the greatest force increase (i.e. peak slope) from the force ski measurement, calculated from the numerical differential of the force curve.

### Data Analysis

Six complete cycles were included in the analysis for each participant and each condition. All data were visually inspected to identify and exclude potential cycles that deviate from the pattern of most cycles. All data were time normalized to 101 points per cycle by linear interpolation. For the analysis, scalars were extracted from the variables (i.e., mean, and peak fascicle and muscle-tendon unit length changes and velocities, and mean normalized and integrated EMG (iEMG) for each participant from the kick phase).

### Statistics

Statistical comparisons of the extracted scalars were performed using Prism (Version 9.1.1, GraphPad Software, LLC, San Diego, CA, USA). All variables were confirmed to be normally distributed using a Shapiro-Wilk’s test (*p* >.05). To assess our hypothesis that there is a stretch-shortening cycle at the muscle-tendon unit level while the muscle fascicles are shortening, we used a single-sided One Sample T-test. For the main variables (MTU and muscle fascicle length and shortening velocity) the hypothetical value was set to 0. For force, the hypothetical value was set to 1 (or 100% of body mass) because participants carry their body mass during the glide phase prior to the kick. Muscles were considered active when EMG activity exceeded two times standard deviation.

To test the effect of speed and incline on the outcome variables a repeated measures One-Way ANOVA with a Geisser-Greenhouse correction was performed. Šídák post-hoc tests were performed in cases of significant main effects of the condition, to compare the effect of incline and speed to DIA_ref_. The significance level was set to *P*<.05 for all statistical tests, and all data are presented as mean ± SD.

## Results

### Cycle characteristics

Repeated measured ANOVA revealed a main effect for average cycle time (*P*<0.0001) and kick duration (*P*<0.0001), while the kick phase was shortest compared to swing and glide phase in all conditions (**Table 1**). Cycle duration was longest for DIA_ref_ and shortest for DIA_steep_. Kick duration was longest for DIA_steep_ for absolute and relative duration. The shortest absolute kick duration was measured for DIA_fast_, while the relative kick duration in DIA_fast_ was not different from DIA_ref_.

**Table 1.** Cycle and phase duration for diagonal stride roller skiing at different incline and speed conditions. N = 13.

| Duration of phases<br>in s (% of cycle) | Cycle | Kick | Swing | Glide |
| --- | --- | --- | --- | --- |
| $DIA_{ref}$ | $1.35 \pm 0.05$<br>(100%) | $0.21 \pm 0.03$<br>(15.9%) | $0.57 \pm 0.04$<br>(42.4%) | $0.59 \pm 0.03$<br>(44%) |
| $DIA_{fast}$ | $1.23 \pm 0.07^*$<br>(100%) | $0.19 \pm 0.04^*$<br>(15.6%) | $0.53 \pm 0.04^{**}$<br>(43.0%) | $0.53 \pm 0.03^{**}$<br>(43.5%) |
| $DIA_{steep}$ | $1.17 \pm 0.05^*$<br>(100%) | $0.24 \pm 0.04^*$<br>(20.8%)** | $0.51 \pm 0.04^{**}$<br>(43.5%) | $0.44 \pm 0.03^{**}$<br>(37.8%)** |

Values are mean ± SD; s = seconds; $DIA_{ref} = 2.5 \text{ m}\cdot\text{s}^{-1}$ , $5^\circ$ ; $DIA_{fast} = 3.5 \text{ m}\cdot\text{s}^{-1}$ , $5^\circ$ ; $DIA_{steep} = 2.5 \text{ m}\cdot\text{s}^{-1}$ , $10^\circ$ ; \* = significantly different from $DIA_{ref}$ ( $P<.05$ ); \*\* = significantly different from $DIA_{ref}$ ( $P<.01$ ).

### Binding forces and joint kinematics

During the kick phase, the ankle joint dorsiflexed in the beginning of the kick and subsequently underwent plantarflexion, while the knee joint underwent a flexion – extension cycle **(Figure 1)**. During the swing phase, forces were zero, while the ankle joint dorsiflexed, and the knee joint flexed. During the glide phase, the force was around the normalized body mass. In all conditions, force declined slightly to 0.8 ± 0.4 N during the glide phase, prior to an increase in force to 1.4 ± 0.6 N before the end of the cycle. During force decline, the ankle dorsiflexed, and the knee extended before they changed direction in the end of the glide phase.

**Figure 1.**
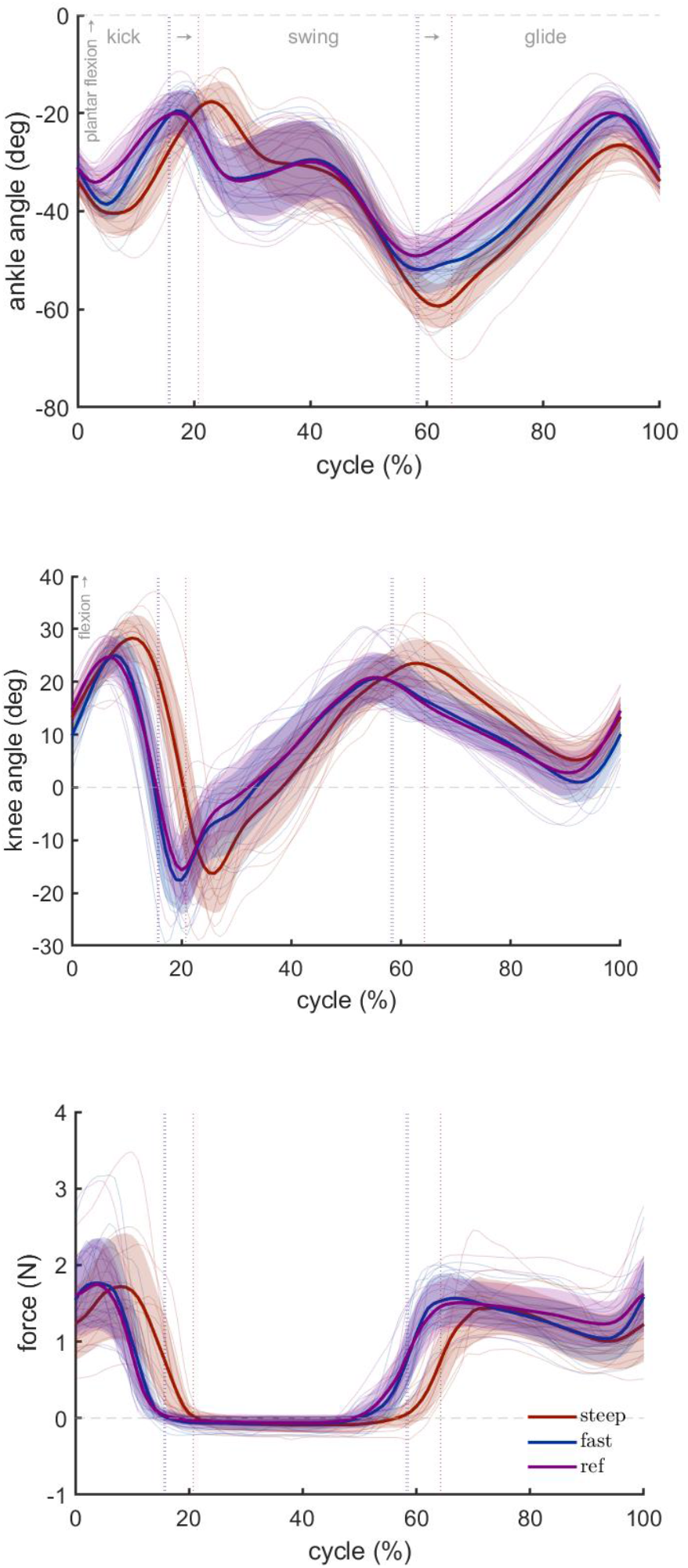
Mean and individual values for ankle joint angle, knee joint angle and force measured under the ski bindings during diagonal style roller skiing during a steep (red), a fast (blue) and a reference condition (ref; purple). Vertical lines denote the phase transition between kick, swing and glide phase for each condition. N = 13.

### Fascicle and MTU behaviour

Gastrocnemius muscle fascicles shortened during the kick phase (9.1 ± 2.7 mm; *P*<0.0001), while the MTU followed a stretch-shortening cycle, with a stretch of 9.9 ± 0.3 mm (*P*<0.0001) and a shortening of 24.3 ± 7.8 mm (*P*<0.0001). Total muscle-tendon unit shortening was 14.4 ± 8.7 mm, which was significantly more than total fascicle shortening (*P*<.0001) (**Figure 2**). Accordingly, peak shortening velocity was significantly slower for fascicles (73.8 ± 32.0 mm·s^−1^) compared to the muscle-tendon unit (375.3 ± 64.8 mm·s^−1^) (**Figure 2** and **Figure 3**). During swing and glide phases, fascicles followed the pattern of the muscle-tendon unit length changes at slower velocities.

**Figure 2.**
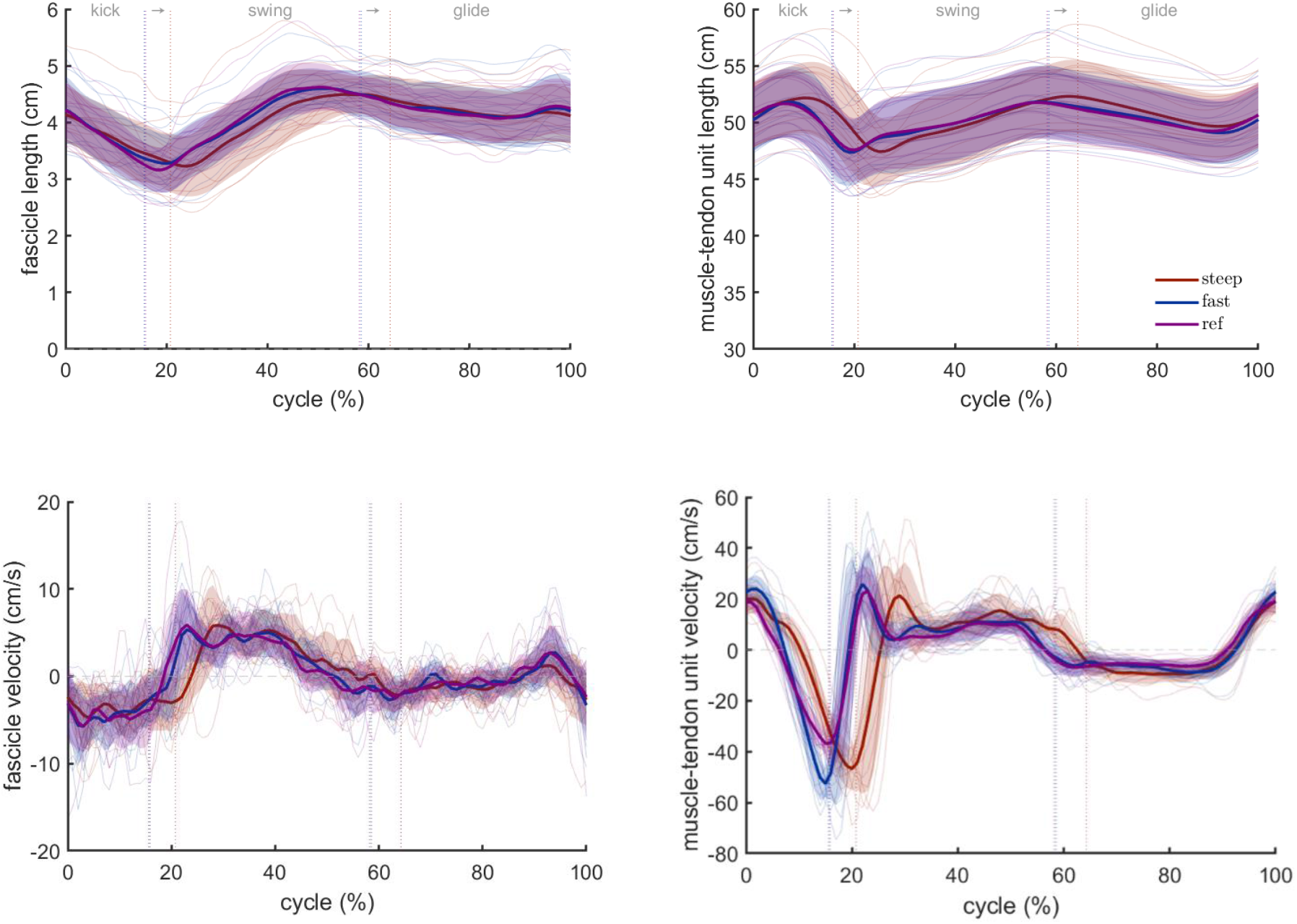
Mean and individual data for fascicle and muscle-tendon unit length and velocity during roller skiing at a steep (red), a fast (blue) and a reference condition (ref; purple). Vertical lines denote the phase transition between kick, swing and glide phase for each condition. N = 13.

**Figure 3.**
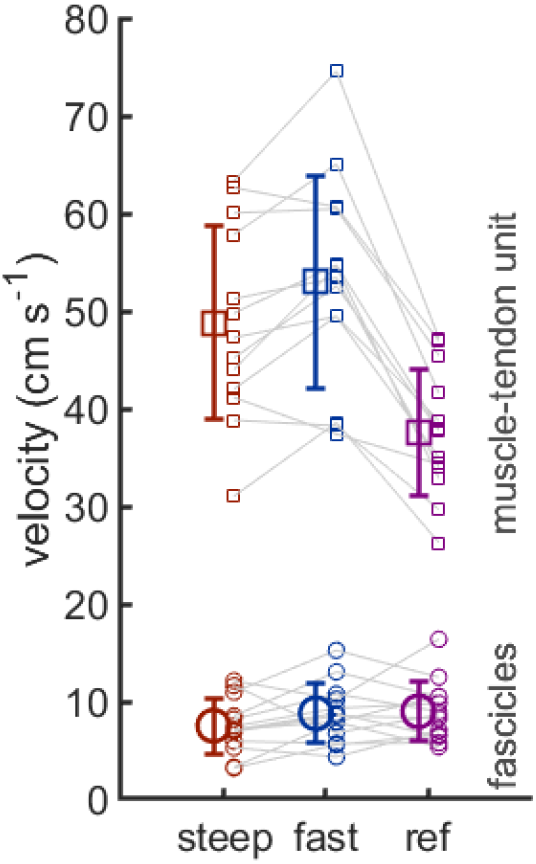
Mean and individual data points for fascicle and muscle-tendon unit peak shortening velocities during the kick phase of diagonal stride roller skiing at a steep (red), a fast (blue) and a reference condition (ref; purple). Shortening is denoted as positive velocities. N = 13.

Across conditions, comparison of operating fascicle length showed no differences (*P*>0.05). However, fascicle length change was significantly smaller in DIA_steep_ (7.6 ± 3.2 mm) compared to DIA_ref_ (9.1 ± 2.7 mm), with a mean difference of 1.5 mm (*P*=0.027). Multiple-comparison tests also showed that mean GM fascicle shortening velocity was slower in DIA_steep_ (34.2 ± 14.8 mm·s^−1^) compared to DIA_ref_ (44.8 mm·s^−1^; *P*=0.0007). Muscle-tendon unit peak length was longer in DIA_steep_ (522.2 ± 31.8 mm) and DIA_fast_ (518.6 ± 31.4 mm) compared to DIA_ref_ (516.7 ± 30.4 mm). Length changes during the stretch phase were 6.5 mm greater for DIA_fast_ (P=0.0003) and 6.1 mm greater for DIA_steep_ (P=0.0015) compared to DIA_ref_. Peak muscle-tendon unit shortening velocity was also greater for both other conditions (DIA_ref_: 375.3 ± 64.8 mm·s^−1^, DIA_fast_ 530.0 ± 109.0 mm·s^−1^, and DIA_steep_ 487.9 ± 98.9 mm·s^−1^), with a mean difference of 154.7 mm·s^−1^ (P<.0001) and 112.6 mm·s^−1^ (P<.0001) for DIA_fast_ and DIA_steep_, respectively (**Figure 3**).

### Muscle activity

Gastrocnemius lateralis and soleus were active during the kick phase (*P*<0.0001) and inactive during the swing phase (**Figure 4**). Both muscles show another increase in activity at the swing to glide transition and the last part of the stride cycle, which was only significant for soleus muscle (*P*=0.0227). At increased speed, peak muscle activity was greater in gastrocnemius (23.1%, *P*=0.0006) and soleus (10.7%, *P*=0.025). At increased incline, peak muscle activity was also greater in gastrocnemius (23.8%, *P*=0.037) and soleus (15.4%, *P*<0.0001). Tibialis anterior muscle was also active during the kick phase but peak activity was during the swing phase.

**Figure 4.**
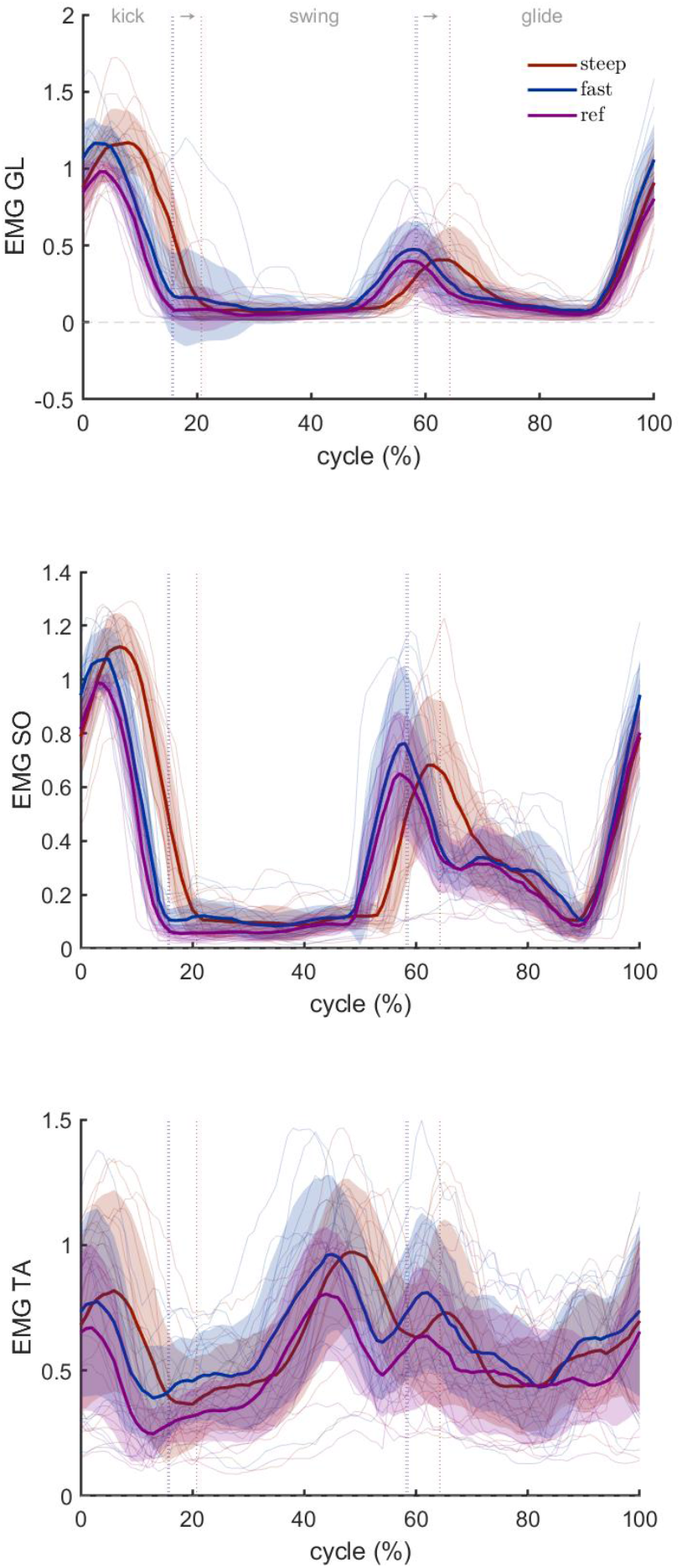
Mean and individual muscle activity (electromyography data) during roller skiing at a steep (red), a fast (blue) and a reference condition (ref; purple). Muscle activity data were normalised to the peak activity during the reference condition. Vertical lines denote the phase transition between kick, swing and glide phase for each condition. N = 13.

## Discussion

Combining ultrasound with motion analysis during diagonal style roller skiing, we found that gastrocnemius muscle fascicles actively shortened during the kick phase, while the muscle-tendon unit underwent a stretch shortening cycle. The muscle-tendon unit operated at significantly higher velocities compared to the fascicles. In the last part of the glide phase, which succeeds the kick, MTU already starts to lengthen, while muscle activity increases, and fascicles contract isometrically (i.e. fascicle velocity is around zero).

As hypothesised, skiers had a longer kick duration at greater incline (absolute and relative), which was accompanied by slower fascicle shortening velocity and higher muscle activity. Speed increase did not result in statistical differences in fascicle velocities or integrated muscle activity, but peak muscle activity was greater with both greater incline and speed. During the kick phase, muscle-tendon unit stretch was larger and faster when speed and incline were greater but muscle-tendon unit shortening velocity, i.e., recoil, was unaffected.

### Uncoupled fascicle and muscle-tendon unit behaviour during the kick

During running elastic elements in the muscle-tendon unit enable energy conservation. The mechanism is characterised by isometric or concentric fascicle work with slow velocities during the stance phase, while the muscle-tendon unit stretches and shortens at much greater velocities (Lichtwark et al., 2007; Roberts and Azizi, 2011). Our data show that muscle fascicles follow a comparable pattern during the kick phase of diagonal style roller skiing as during running, despite the glide phase prior to the kick during diagonal stride skiing. Similar mechanisms during skiing and running may be surprising considering that ground contact duration during skiing is much longer due to the added glide phase. Similar to walking, the foot’s duty factor (i.e., percent of cycle time with foot contact) during skiing is greater than 0.5. Notably, race walking was characterised by significantly less energy efficient contractile patterns compared to running (Cronin et al., 2016). Thus, the glide phase during diagonal style skiing, where calf muscles are mostly passive, seems to be unlike ground contact phases during walking and race walking, where muscle activation is high during the whole ground contact phase. Our result support, at least in part, the description of diagonal stride skiing as “grounded running” by Pellegrini et al. (2014). The term originally describes a certain type of animal locomotion, highlighting the combination of characteristics from running where the gliding phase replaces parts of the aerial phase. Notably, grounded running has been associated with reduced musculoskeletal loading compared to other gait (Bonnaerens et al., 2019).

Conservation of elastic energy and/or power amplification - as apparent in different types of locomotion (Roberts and Azizi, 2011) - may also play a role during diagonal style skiing. Komi and Norman (1987) were the first to demonstrate a stretch-shortening pattern of the muscle-tendon unit during the kick phase of this gait. Interestingly, the stretch started already at the end of the glide phase, immediately before the kick, and was suggested to be used as preloading phase. Our data shows a similar pattern in addition to an increase in gastrocnemius and soleus muscle activity (Figure 4) in the end of the glide phase, supporting the notion that the muscles are active. This stretch-shortening cycle might indicate elastic energy storage and release during the glide- and kick-phases. Based on fluctuations of mechanical energy, Kehler et al. (2014) concluded that most of the mechanical energy is dissipated during the glide phase (through ski/snow friction) rather than being stored elastically. Hence, unlike running, the elastic energy released during the kick-phase of diagonal stride skiing is most likely actively generated during a brief preload-phase prior to the kick, rather than a result of mechanical energy conservation throughout the stride cycle (as in a bouncing gait). The notion of energy dissipation during the glide phase based on mechanical energy transfer is consistent with our data showing little muscle activation during this phase (e.g. in Brennan et al., 2018; Kehler et al., 2014).

Jumping is another movement with similarities to incline skiing because both require positive work (Dahl et al., 2017). During jumping, elastic mechanisms in the muscle-tendon unit allow for power amplification by stretching at slower rate than recoiling (Farris et al., 2016; Wade et al., 2018). Accordingly, the lengthening of the muscle-tendon unit with concurrent absence of fascicle lengthening in the end of the glide phase indicates a power amplification mechanism similar to jumping. However, measurements of tendon and other elastic tissue velocities are required to conclude about the contribution of power amplification during skiing. Besides jumping, positive work production of fascicles has been shown during running at incline (Lichtwark and Wilson, 2006) and during accelerative movements such as accelerative walking (Farris and Raiteri, 2017) and sprint acceleration (Werkhausen et al., 2021), supporting the hypothesis that the compliant lower leg muscles contribute to positive work production when required. Indeed, the average fascicle shortening during DIA_ref_ of 9.1 mm was similar to fascicle shortening during accelerative sprinting (9.6 mm) and also muscle-tendon unit velocities were similar with around 50 cm·s^−1^ (Werkhausen et al., 2021).

Overall, the simultaneous muscle-tendon unit stretch and fascicle shortening in the end of the glide phase and the beginning of the kick phase indicates that using stretch shortening cycle exercises may be beneficial in training for skiers. They may also benefit from adjusting their technique for optimal energy storage by preloading elastic tissues in the end of the glide phase.

### Speed and incline affect fascicle and muscle-tendon dynamics

Increase in speed and incline altered cycle characteristics, forces and kinematics during diagonal style roller skiing, in line with previous studies (Nilsson et al., 2004; Pellegrini et al., 2013). These differences were associated with differences in fascicle and muscle-tendon unit behaviour. At increased speed, greater muscle activation for gastrocnemius and soleus muscles may be driven by the shorter kick phase. Fascicle behaviour, such as operating length and velocity, was preserved although peak muscle-tendon unit length was longer and velocity was greater in DIA_fast_ compared to DIA_ref_. Interestingly, this modulation of muscle behaviour to speed increases resembles that of running at increasing speeds (Farris and Sawicki, 2012; Werkhausen et al., 2019). Based on the modulation of further speed increases during running, we speculate that greater increases in skiing speed (than in the current protocol) would lead to higher fascicle shortening velocities. Nevertheless, concomitant increase in muscle-tendon unit velocity (while fascicle velocity did not differ) may indicate a greater stretch of elastic elements and thereby greater potential for energy recycling and power amplification when increasing speed with 1 m·s^−1^ or 40%.

At greater incline, when the resistive force was higher, skiers increased absolute and relative kick duration. Contrary to speed increases, fascicle velocity was slower at DIA_steep_, although muscle-tendon unit excursion and velocity were greater, likely allowing for increased force potential (according to the force-velocity relationship) of the muscle and thereby lower energy requirements (Lieber and Fridén, 2000). Yet, the higher peak muscle activity counteracts a possible energy saving of slower fascicle operating velocities by requiring more energy. Besides fascicle velocity, operating lengths have a potential effect on energy consumption (Beck et al., 2022) but were not present when comparing our conditions. Lichtwark and Wilson (2006) reported that fascicles operated at longer length when running uphill compared to level running but concluded that the small difference was marginal. Overall, differences in muscle behaviour in the different speed and incline conditions indicate that training at specific conditions may be necessary if athletes wish to train specific contractile modes.

### Methodological considerations

Roller skiing on a treadmill closely resembles skiing on snow but it is important to note that small differences may influence the results on muscle behaviour found in our study.

Myklebust et al. (2022) found differences in pole push time and hip kinematics during skating (V2) and several studies have discussed the importance of grip on snow for diagonal stride (e.g. Kehler et al., 2014). In our study, we used very experienced skiers and placed the ratchet mechanism on the front wheel to reduce grip, to minimise the differences between roller skiing and on-snow skiing. We consider including both sexes in our study a strength in general, but it is important to note that we have not corrected for existing sex differences (Stöggl et al., 2018).

### Conclusion

Uncoupling of fascicle behaviour of the gastrocnemius muscle from the limb joint movement during diagonal style cross-country skiing suggests the use of elastic energy at different inclines and speeds. Skiers may use the end of the glide phase to preload the muscle-tendon unit for the stretch shortening cycle during the kick, where muscle fascicles shorten actively. The contractile pattern was overall similar when varying speed or incline. Both increased incline and speed resulted in increased muscle activation, whereas greater incline additionally enabled slower shortening velocities.

## Competing interests

No competing interests declared.

## Funding

This research received no specific grant from any funding agency in the public, commercial or not-for-profit sectors.

## Data availability

The data that support the findings of this study are available from the corresponding author [AW] upon reasonable request.

## Notes

### Competing Interest Statement

The authors have declared no competing interest.

